# Intermittent theta burst stimulation (iTBS)-induced changes of resting-state brain entropy (BEN)

**DOI:** 10.1101/2024.05.15.591015

**Authors:** Pan-Shi Liu, Dong-Hui Song, Xin-Ping Deng, Yuan-Qi Shang, Qiu Ge, Ze Wang, Hui Zhang

## Abstract

Intermittent theta burst stimulation (iTBS) is a novel protocol of repetitive transcranial magnetic stimulation (rTMS). While iTBS has shown better therapeutic effects for depression than conventional high-frequency rTMS (HF-rTMS), its underlying neuronal mechanism remains elusive. Brain entropy (BEN), a measure of irregularity of brain activity, has recently emerged as a novel marker of regional brain activity. Our previous studies have shown the sensitivity of BEN to depression and HF-rTMS, suggesting BEN as a sensitive tool for understanding the brain mechanism of iTBS. To assess this possibility, we calculated BEN using resting state fMRI data provided by an open dataset in OpenNeuro. Sixteen healthy participants underwent 600 pulses of iTBS applied over the left dorsolateral prefrontal cortex (L-DLPFC) at two intensities (90% and 120% of individual resting motor threshold (rMT)) on separate days. We assessed the pre-post stimulation BEN difference and its associations with neurotransmitter receptor and transporter binding maps. Our results showed that subthreshold iTBS (90% rMT) decreased striatal BEN, while suprathreshold iTBS (120% rMT) increased striatal BEN. We also found significant differences in the spatial correlation between BEN changes induced by different stimulation intensities and various neurotransmitters. These results suggest that differences in BEN caused by iTBS stimulation intensity may be related to the release of other neurotransmitters. The study underscores the significance of iTBS stimulation intensity and provides a basis for future clinical investigations to identify stimulation intensities with good therapeutic benefits.

## 1 Introduction

Non-invasive brain stimulation (NIBS) offers a safe way to investigate the causal relationship between brain function and behavior and subsequently represents one of the most significant advancements in neuroscience (Miniussi, Harris et al. 2013, Polanía, Nitsche et al. 2018). NIBS not only provides a valuable approach to exploring the neural mechanisms underlying human behavior but also holds significant potential for various clinical applications (Hummel and Cohen 2006, Schulz, Gerloff et al. 2013). Repetitive transcranial magnetic stimulation (rTMS) is among the most widely used NIBS techniques adopted in both basic neuroscience research and clinical applications, especially in neuropsychological and neuropsychiatric disorders (Machii, Cohen et al. 2006, Ridding and Rothwell 2007, Bakker, Shahab et al. 2015, Somaa, de Graaf et al. 2022). An increasing number of studies have shown that rTMS can induce neuroplasticity across brain regions, systems, and timescales (Song, Chang et al. 2019, Shang, Chang et al. 2020, Feng, Deng et al. 2022, Fitzsimmons, Oostra et al. 2023, Deng, Chen et al. 2024), through the long-term potentiation/depression (LTP/LTD) of excitatory synaptic transmission (Esser, Huber et al. 2006, Hoogendam, Ramakers et al. 2010). Conventional high-frequency rTMS (HF-rTMS >5 Hz) over the left dorsolateral prefrontal cortex (L-DLPFC) can increase executive control functions such as working memory and is a widely adopted treatment protocol for psychiatry disorders such as major depression disorder (MDD) (Avery, Holtzheimer III et al. 2006, Teng, Guo et al. 2017) and substance use disorders (SUD) (Gorelick, Zangen et al. 2014, Coles, Kozak et al. 2018). Intermittent theta burst stimulation (iTBS) (Rounis and Huang 2020) is a new rTMS protocol motivated by the brain’s natural theta rhythm occurring in the hippocampus (Klomjai, Katz et al. 2015). When applied to the L-DLPFC, iTBS shows similar efficacy, acceptability, and safety to HF-rTMS for MDD treatment with even marginally better treatment effects than HF-rTMS regarding remission rate (Bulteau, Sébille et al. 2017, Kishi, Sakuma et al. 2023). Moreover, iTBS consists of short trains of stimuli played at high frequency, which offers a key advantage over conventional HF-rTMS in terms of shorter stimulation duration (Chung, Hoy et al. 2015, Mendlowitz, Shanbour et al. 2019, Voigt, Leuchter et al. 2021). Several studies have demonstrated that variations in iTBS intensities produce different effects in both healthy and diseased individuals (Chung, Rogasch et al. 2018, Lee, Corlier et al. 2021, Zhang, Kan et al. 2022) that might account for the heterogeneous results (de Boer, Schluter et al. 2021, Pabst, Proksch et al. 2022). While these iTBS results are encouraging, their underlying neuromechanisms are unknown (Pabst, Proksch et al. 2022). We have previously studied rTMS brain effects using resting state fMRI-derived regional brain entropy (BEN) (Donghui Song 2017, Song, Chang et al. 2019). In this study, we aimed to identify iTBS-induced regional BEN alterations and their associations with distributions of major neurotransmitters.

BEN is a robust measure of the human brain activities in normal or disease conditions by depicting the intricate dynamic state of the brain from information theory (Wang, Li et al. 2014) that provides complementary information to other measures of spontaneous brain activity, such as the fractional amplitude of low-frequency fluctuation (fALFF) (Zou, Zhu et al. 2008) and cerebral blood flow (Wang, Aguirre et al. 2008, Zou, Wu et al. 2009, Li, Zhu et al. 2012, Song, Chang et al. 2019). Studies of regional BEN based on resting-state functional magnetic resonance imaging (rs-fMRI) have been growing rapidly in recent years (Wang, Li et al. 2014, Wang 2021, Song, Jann et al. 2024). The relationship between BEN and cognitive function has been identified (Wang, Li et al. 2014, Wang 2021, Lin, Chang et al. 2022, Del Mauro and Wang 2023), and mental disorders related to BEN patterns have also been found in a variety of brain diseases (Liu, Song et al. 2020, Wang and Initiative 2020, Jiang, Cai et al. 2023), including MDD (Liu, Song et al. 2020, Dong-Hui Song 2024). BEN also reflects the effects of pharmacological (Chang, Song et al. 2018, Liu, Song et al. 2020) and nonpharmacological treatment for depression (Dong-Hui Song 2024). More importantly, we found that BEN in the medial orbitofrontal cortex and subgenual anterior cingulate cortex (MOFC/sgACC), which are associated with emotion regulation, was reduced by HF-rTMS over L-DLPFC in normal young adults (Donghui Song, Zhang et al. 2017, Song, Chang et al. 2019). HF-rTMS over L-DLPFC is also used to intervene in SUD (Li, Hartwell et al. 2013, Lefaucheur, Aleman et al. 2020) and our recent study found that participants with SUD had higher BEN in the prefrontal and mesolimbic relative to healthy controls (Jiang, Cai et al. 2023). A recent finding from Timothy Jordan et al. (2024) revealed that individuals who smoke exhibited a higher baseline BEN compared to healthy controls, consistent with our previous findings (Li, Fang et al. 2016, Jiang, Cai et al. 2023). Furthermore, they found that HF-rTMS over L-DLPFC reduced BEN in the insula and DLPFC in SUD, and reduced self-reported cigarette craving (Jordan, Apostol et al. 2024). These studies all indicate that BEN is sensitive to the effects of rTMS in both healthy and diseased conditions.

As previously mentioned, synaptic LTP/LTD are widely regarded as fundamental mechanisms underlying learning and memory (Lynch 2004, Di Filippo, Picconi et al. 2009, Mancini, de Iure et al. 2022), and the neuroplasticity mechanisms of TMS are often attributed to the induction of LTP/LTD (Esser, Huber et al. 2006, Hoogendam, Ramakers et al. 2010, Huang, Rothwell et al. 2011). At the cellular level, substantial evidence suggests that dopamine (DA) and acetylcholine (ACh) primarily modulate the magnitude of LTP (Gu 2002, Monday, Younts et al. 2018), while 5-Hydroxytryptamine (5HT), glutamate (Glu) and gamma-aminobutyric acid (GABA) influence LTD (Massey and Bashir 2007, Lovinger 2010, Monday, Younts et al. 2018). Results from animal experiments have already demonstrated that rTMS increases the release of DA and Glu in the striatum (Keck, Welt et al. 2002, Zangen and Hyodo 2002) and also affects other neurotransmitters (Albert, Cook et al. 2009). In a clinical context, research on the therapeutic mechanisms of rTMS has also clearly demonstrated its effects on neurotransmitters and GABA and Glu neurotransmitter systems are central to the pathophysiology of depression and are potential targets of HF-rTMS over L-DLPFC (Post and Keck 2001, Chervyakov, Chernyavsky et al. 2015, Dubin, Mao et al. 2016). A recent review showed that rTMS and MRS (magnetic resonance spectroscopy) reveal complementary and comprehensive information on neurotransmission, both in normal adults and in disease conditions (Cuypers and Marsman 2021). Specifically regarding iTBS, a review from Larson et al (2015) indicated that repeated theta-burst stimulation induces maximal LTP, primarily because this frequency disables feedforward inhibition and allows sufficient postsynaptic depolarization, and this disinhibitory process involves presynaptic GABA autoreceptors that inhibit GABA release (Larson and Munkácsy 2015). Another review of findings on TBS from animal and human studies suggests that GABA and Glu are critically involved in these mechanisms (Li, Huang et al. 2019). Since iTBS can cause changes in neurotransmitters, theoretically the spatial pattern of BEN changes caused by iTBS should be related to the spatial distribution of neurotransmitters.

Recently, a study by Alkhasli at al. (2019) indicated that different stimulation intensities influence functional connectivity (FC) among the frontal-striatal network (Alkhasli, Sakreida et al. 2019). Following this study using the same dataset, we employ BEN as a reliable index to investigate the mechanism of the iTBS effect on local brain activity. Recent data-sharing efforts allow us to relate iTBS-induced BEN patterns to neurotransmitter receptor and transporter binding maps obtained in PET (Hansen, Shafiei et al. 2022). The present study aims to advance our understanding of the neural mechanisms underlying iTBS by addressing the following three questions: a) whether BEN exhibits sensitivity to the effects of iTBS comparable to that observed with HF-rTMS targeting the L-DLPFC; b) whether BEN can effectively capture the effects of iTBS at varying intensities; and c) the correlation between iTBS-induced BEN patterns and neurotransmitters distribution. Drawing from results and insights from Alkhasli et al. (2019) in the frontal-striatal network, we hypothesize that BEN exhibits differential responses to varying stimulation intensities, primarily in the L-DLPFC and striatum. Additionally, we propose that BEN patterns caused by different iTBS intensities may be related to different neurotransmitters.

## 2 Methods

### 2.1 Datasets

The data of the study from OpenNeuro ds001832 that released by Alkhasli et al. (https://openneuro.org/datasets/ds001832/versions/1.0.1). Sixteen healthy participants were recruited in this dataset (mean age = 27.63, standard deviation (SD)= 6.95; 8 males).

#### 2.1.1 Experimental Procedure

Each participant completes this study over three days. On day 1, they underwent pre-screening, and information consent was completed. On day 2, the resting motor threshold (rMT) was determined using a standardized protocol (Rossi, Hallett et al. 2009, Rossini, Burke et al. 2015). After a 20-minute break, a 10-minute rs-fMRI was conducted. Subsequently, for iTBS measurement, the participants were moved outside the scanner room, and a 3-minute 20-second iTBS over L-DLPFC was applied at either 90% or 120% of the rMT. The order of stimulation intensities (90% and 120%) was counterbalanced and alternated between participants. A 7-minute break after iTBS and another 10-minute rs-fMRI were performed. Participants were instructed to keep their eyes open during rs-fMRI. The measurements on day 3 mirror those of day 2, with the only difference being the change in stimulation intensity.

#### 2.1.2 Magnetic Resonance Imaging

Magnetom Prisma 3.0 T whole-body scanner (Siemens Medical Solutions, Erlangen, Germany) was used to acquire anatomical images and rs-fMRI images.

Anatomical images were acquired using a three-dimensional magnetization-prepared, rapid acquisition gradient-echo sequence (MPRAGE) with the following parameter: 300 repetitions, TR = 2,300 ms, TE = 2.98 ms, 9° flip angle, FOV = 256 mm, 176 sagittal slices, slice thickness = 1 mm and in-plane resolution = 1 × 1 × 1 mm.

Rs-fMRI images were acquired with a gradient echo planar imaging (EPI) sequence with the following parameters: TR = 2,000 ms, TE = 28 ms, 77° flip angle, FOV = 192 mm, 34 axial slices (interleaved acquisition), 3 mm slice thickness. A total of 300 volumes were acquired during 10 min.

#### 2.1.3 iTBS

An anatomical image was used to locate the hand area of the left motor cortex to determine the individual rMT. The head was registered to individual anatomical images using neuronavigation software (TMS navigator, Localite GmbH, Sankt Augustin, Germany). The presumed hand area was identified visually through anatomical landmarks and stimulated with biphasic single pulses using a figure-of-eight coil (MagVenture C-B60) connected to a MagPro stimulator (X100 MagVenture, Farum, Denmark). Electrodes were fitted to the participant’s right index finger and motor evoked potentials were monitored. Stimulation intensity was first increased in 2% steps until the hand area could be determined through a clear supra-threshold (>50 μV) motor-evoked potential. The intensity was then reduced stepwise to find the lowest intensity to still induce a supra-threshold motor-evoked potential.

A total of 600 pulses spaced out over 3 min and 20 s in iTBS protocol that was comprised of 20 trains and 10 theta-bursts and was applied over the L-DLPFC. The mean stimulation sites were x,y,z = (−41,37,31) in Talairach coordinates. The mean subthreshold stimulation applied was 38% and the mean suprathreshold was 50% of the maximum stimulator output.

For more detailed participants information, experimental procedure, MRI acquisition parameters, and iTBS parameters can be found in the original article for the dataset (Alkhasli, Sakreida et al. 2019).

### 2.2 MRI preprocessing

The MRIs were preprocessing using fmriprep (Esteban, Markiewicz et al. 2019) (ver sion=23.1.4) (https://pypi.org/project/fmriprep-docker/), which is based on Nipype (version=1.8.6) (Gorgolewski, Burns et al. 2011), is containerized to docker (version=24.0.7) (https://www.docker.com/) under Ubuntu (version=22.04.3 LTS) (https://releases.ubuntu.com/jammy/), then denoising using based-Nilearn (version=0.10.2) (https://nilearn.github.io/stable/index.html) customed python (version=3.10) (https://www.python.org/) scripts.

#### 2.2.1 Structure MRI preprocessing

Each T1w image was corrected for intensity non-uniformity (INU) with N4BiasFieldCorrection (Tustison, Avants et al. 2010), distributed with ANTs (Avants, Tustison et al. 2009), and used as T1w-reference throughout the workflow. The T1w-reference was then skull-stripped with a Nipype (Gorgolewski, Burns et al. 2011) implementation of the antsBrainExtraction.sh workflow, using OASIS30ANTs as a target template. Brain tissue segmentation of cerebrospinal fluid (CSF), white matter (WM), and gray matter (GM) was performed on the brain-extracted T1w using fast (Zhang, Brady et al. 2001). Volume-based spatial normalization to MNI152NLin2009cAsym was performed through nonlinear registration with antsRegistration, using brain-extracted versions of both the T1w reference and the T1w template. The ICBM 152 Nonlinear Asymmetrical template version 2009c (Fonov, Evans et al. 2009) was selected for spatial normalization and accessed with TemplateFlow (Ciric, Thompson et al. 2022).

#### 2.2.2 fMRI preprocessing

First, a reference volume and its skull-stripped version were generated using a custom methodology of fMRIPrep. Head-motion parameters with respect to the BOLD reference (transformation matrices, and six corresponding rotation and translation parameters) are estimated before any spatiotemporal filtering using mcflirt (Jenkinson, Bannister et al. 2002). BOLD runs were slice-time corrected to 0.96s (0.5 of slice acquisition range 0s-1.92s) using 3dTshift from AFNI (Cox and Hyde 1997). The BOLD time-series (including slice-timing correction) were resampled onto their original, native space by applying the transforms to correct for head motion. These resampled BOLD time series will be referred to as preprocessed BOLD in the original space. The BOLD reference was then co-registered to the T1w reference using mri_coreg followed by flirt (Jenkinson and Smith 2001) with the boundary-based registration (Greve and Fischl 2009) cost-function. Co-registration was configured with six degrees of freedom. Several confounding time series were calculated based on the preprocessed BOLD: framewise displacement (FD), DVARS, and three region-wise global signals. FD was computed using two formulations following Power (absolute sum of relative motions,(Power, Barnes et al. 2012)) and Jenkinson (relative root mean square displacement between affines,(Jenkinson, Bannister et al. 2002)). FD and DVARS are calculated both using their implementations in Nipype (Gorgolewski, Burns et al. 2011). The three global signals are extracted within the CSF, the WM, and the whole-brain masks. Additionally, a set of physiological regressors was extracted to allow for component-based noise correction (Behzadi, Restom et al. 2007). Principal components are estimated after high pass filtering the preprocessed BOLD time series (using a discrete cosine filter with 128s cut-off) for the two CompCor variants: temporal (tCompCor) and anatomical (aCompCor). The BOLD time series were resampled into MNI152NLin2009cAsym spaces. First, a reference volume and its skull-stripped version were generated using a custom methodology of fMRIPrep. All resamplings can be performed with a single interpolation step by composing all the pertinent transformations (head-motion transform matrices, susceptibility distortion correction when available, and co-registrations to anatomical and output spaces). Gridded (volumetric) resamplings were performed using antsApplyTransforms (ANTs), configured with Lanczos interpolation to minimize the smoothing effects of other kernels (Lanczos 1964). The images preprocessed by fmriprep are denoised using the based on Nilearn customed python scripts, including linear trend removal, bandpass filter (0.009-0.08), regression out confounds including WM, CSF signal, six head movement parameters, FD, std_dvars, rmsd, tcompcor and first three componence of c_comp_cor, then smoothing with an isotropic Gaussian kernel (FWHMLJ=LJ6 mm), while mean FD were calculated from FD for each participant.

### 2.4 BEN mapping

The voxel-wise BEN maps were calculated from the preprocessed rs-fMRI using the BEN mapping toolbox (BENtbx) (Wang, Li et al. 2014) based on sample entropy (Richman and Moorman 2000). The toolbox can be found at https://www.cfn.upenn.edu/zewang/BENtbx.php and https://github.com/zewangnew/BENtbx. The window length was set to 3 and the cutoff threshold was set to 0.6 according to our experimental optimization (Wang, Li et al. 2014). The first four volumes were discarded for signal stability and to reduce variability due to noise, BEN maps were smoothed with an isot ropic Gaussian kernel with FWHMLJ=LJ8 mm. These parameter settings are consistent with our previous studies, and more details of BEN calculation can be found in our previous studies (Wang, Li et al. 2014, Song, Chang et al. 2019, Song, Chang et al. 2019, Lin, Chang et al. 2022).

### 2.5 Statistical analysis

#### 2.5.1 Interactive effects between stimulation intensity and time point

A 2×2 repeated measures analysis of variance (ANOVA) was used to evaluate th e interaction between stimulation intensity (90% vs. 120%) and time point (Pre vs. Po st), this analysis was performed using Statistical Parametric Mapping (SPM12, WELL COME TRUST CENTRE FOR NEUROIMAGING, London, UK, http://www.fil.ion.ucl.ac.uk/spm/software/spm12/) (Friston, Holmes et al. 1994). The significance level was defined as p < 0.05 after multiple comparison corrections (voxel-wise p < 0.005, cluster size > 29) using AFNI 3dClustSim (Cox 1996). For multiple comparison correction, AFNI 3dFWHMx was employed to estimate the autocorrelation function (ACF) using residual maps obtained from each participant’s BEN maps output by SPM12, then a pre-defined region of interest (ROI) including the L-DLPFC (Brodmann areas 46, 9) a nd striatum was utilized to correct the F statistical maps using 3dClustSim. Predefined masks for the L-DLPFC and striatum are available from https://neurovault.org/collections/QYIWAEYS/.

#### 2.5.2 Effects of different stimulation intensities on BEN

The average BEN values of the clusters exceeding the threshold from ANOVA were extracted, and then customized Python scripts were used to examine the changes in BEN before and after stimulation using the paired T-test for different stimulation intensities.

#### 2.5.3 The relationship between iTBS-induced BEN patterns and neurotransmitters

Neurotransmitter maps were acquired from https://github.com/netneurolab/hansen_receptors/tree/main/data/PET_nifti_images (Hansen, Shafiei et al. 2022). The 39 neurotransmitter maps were generated from PET tracer binding images from 19 different neurotransmitter receptors, transporters, and receptor-binding sites across nine different neurotransmitter systems (dopamine, norepinephrine, serotonin, acetylcholine, glutamate, GAB A, histamine, cannabinoid, and opioid) (Hansen, Shafiei et al. 2022). SPM12 was used to calculate the voxel-wise BEN differences before and after stimulation using the paired T-test. T-statistical maps and neurotransmitter maps were resampled to the same resolution (3 mm × 3 mm × 3 mm). Pearson correlation between the stimulation-induced (Post-Pre) BEN changes and the neurotransmitter distribution of each neurotransmitter map was calculated. Given that a very small correlation coefficient can lead to ver y significant results on a large amount of data (49233 voxels), we selected the correl ation coefficient r>0.1 (p<1×10^^-119^) as significant results. The correlation coefficients were subsequently converted into z-scores, and then compared to assess significant dif ferences between them, significance level was defined as z-score>10, p << 2.56×10^-5^ (Bonferroni correction, 0.001/39).

## 3 Results

### 3.1 Interactive effects between stimulation intensity and time in the striatum

ANOVA revealed significant interactive effects on BEN between stimulation intensities (90% vs 120%) and time points (Pre vs Post) in the striatum, encompassing the bilateral putamen and right caudate (see Fig 1, Table 1).

**Fig 1.**
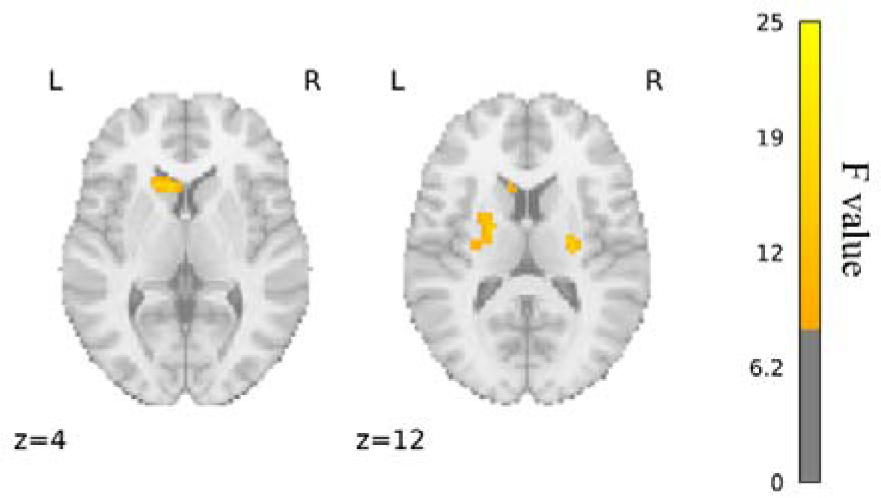
Interaction stimulation intensity and time point. The number beneath each slice indicates its location along the z-axis in the MNI space, and L means left hemisphere, R means right hemisphere, and colorbar indicates F values.

**Table 1.**
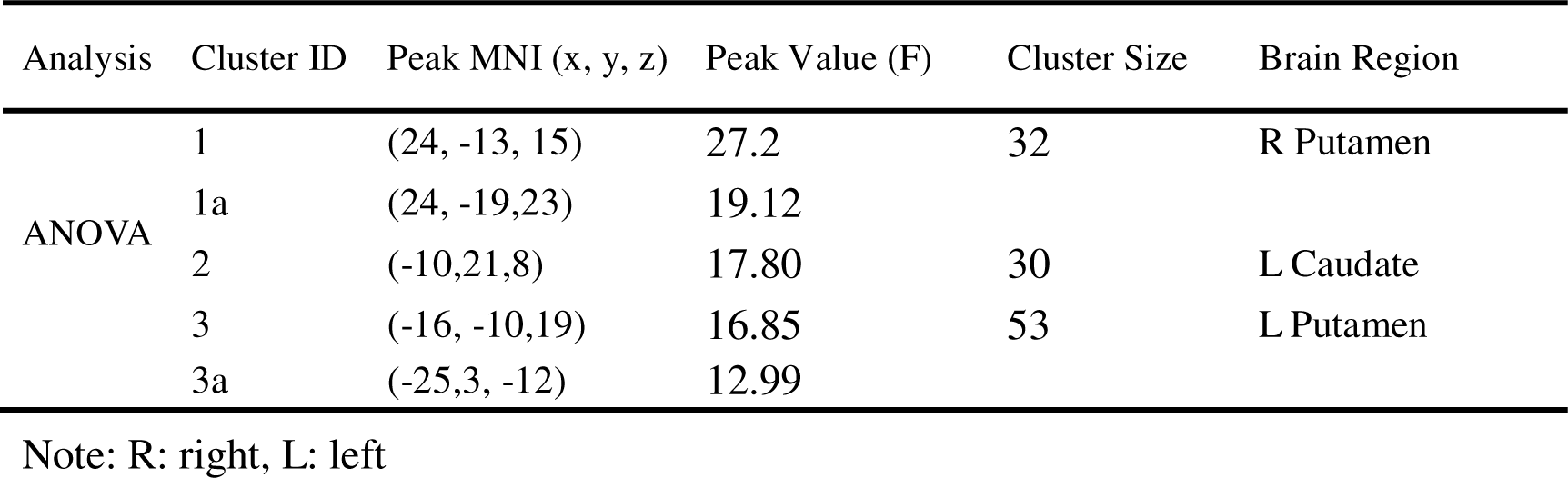
Clusters size table for ANOVA.

| Analysis | Cluster ID | Peak MNI (x, y, z) | Peak Value (F) | Cluster Size | Brain Region |
| --- | --- | --- | --- | --- | --- |
| ANOVA | 1 | (24, -13, 15) | 27.2 | 32 | R Putamen |
|  | 1a | (24, -19, 23) | 19.12 |  |  |
|  | 2 | (-10, 21, 8) | 17.80 | 30 | L Caudate |
|  | 3 | (-16, -10, 19) | 16.85 | 53 | L Putamen |
|  | 3a | (-25, 3, -12) | 12.99 |  |  |

Note: R: right, L: left

**Table 2.**
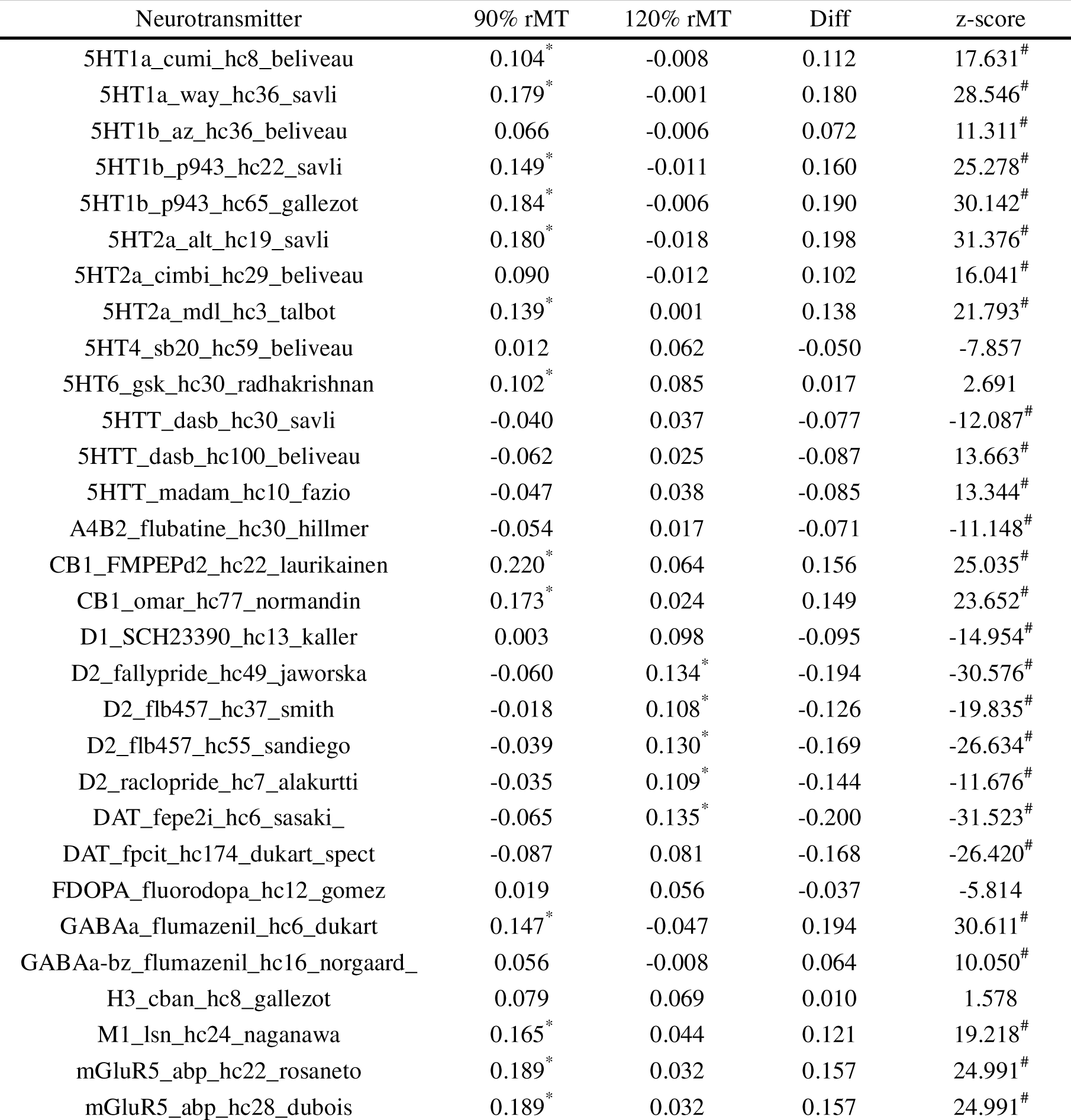
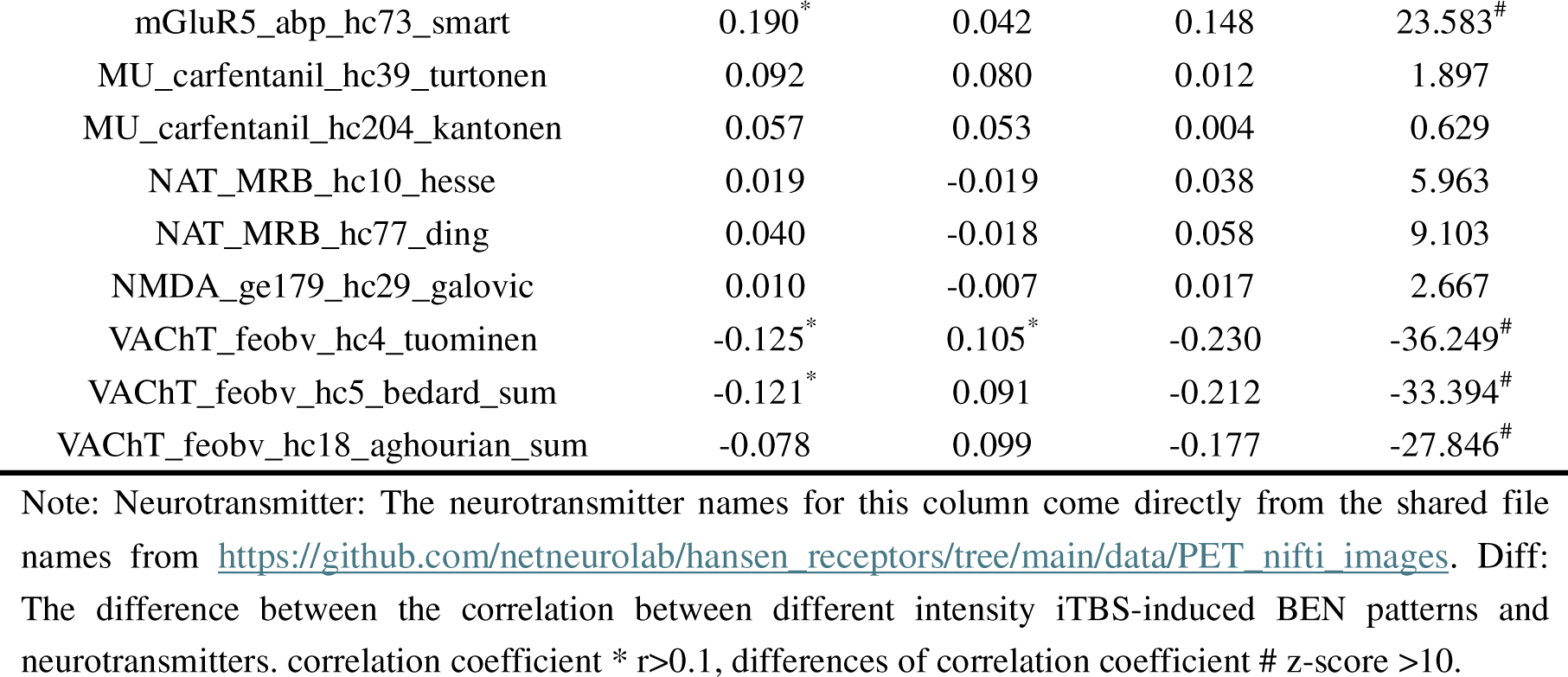
The correlation coefficient of iTBS-induced BEN patterns and neurotransmitter.

### 3.2 Effects of different stimulation intensities on BEN

The average BEN values were calculated for each cluster. Like our voxel-wise level findings, we observed a significant interaction between stimulation intensity and time point for the average BEN within each cluster. Subthreshold iTBS led to a decrease in BEN across all clusters, while suprathreshold iTBS resulted in an increase in BEN across all clusters. These results were also evident at the individual level (see Fig 2).

**Fig 2.**
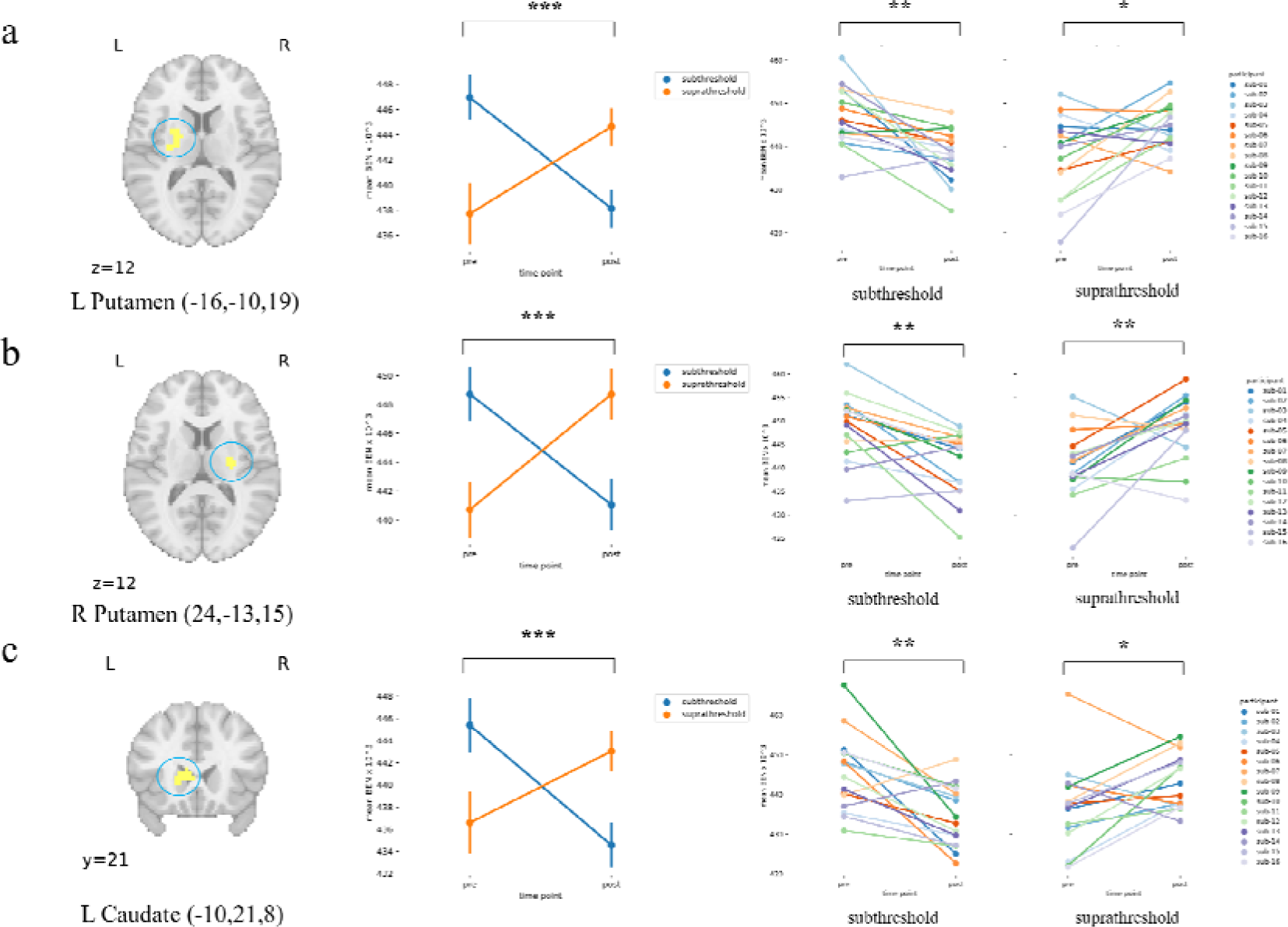
Effects of different stimulation intensities on BEN for group and individual. The changes of BEN before and after stimulation with different stimulation intensities in **a)** the left putamen, **b)** the right putamen, and c) the left caudate. Asterisks indicate significance levels for 2×2 repeated measures ANOVA or paired samples T-test, * p<0.05, ** p<0.01, *** p<0.001. Left: The location of clusters on the anatomical template (the blue circle), the numbers in brackets are peak MNI coordinates in the clusters, the number beneath each slice indicates its location along the z-axis or y-axis in the MNI space, and L means left hemisphere, R means right hemisphere. Meddle: group means changes in BEN before and after stimulation with different stimulation intensities, y-axis represents the mean BEN values × 10^^3^, while the x-axis corresponds to different time points (Pre vs Post), yellow represents suprathreshold (120% rMT) iTBS and blue represents subthreshold (90% rMT) iTBS. Line height represents the standard error (SE), and maker represents the mean BEN values 10^^3^. Right: Mean BEN changes before and after stimulation for each subject for different stimulation intensities, the y-axis represents the mean BEN values × 10^^3^, while the x-axis corresponds to different time points.

### 3.3 The relationship between iTBS-induced BEN patterns and ne urotransmitters

The spatial analysis of BEN patterns induced by iTBS and neurotransmitters showed different patterns depending on the stimulation intensity. For subthreshold iTBS, there were significant positive correlations with 5HT, cannabinoid receptor 1 (CB1), GABA, muscarinic acetylcholine receptor M1 (M1), and metabotropic glutamate receptor 5 (mGluR5), and significant negative correlations with vesicular acetylcholine transporter (VAChT). On the other hand, suprathreshold iTBS resulted in BEN change patterns positively correlated with dopamine receptor D2 (D2), dopamine transporter (DAT), and VAChT, with no significant negative correlations identified. Importantly, the relationship between BEN change patterns and neurotransmitters differed significantly between subthreshold and suprathreshold iTBS for these neurotransmitters (see Table 3).

## 4 Discussion

Our research findings show that different intensities of iTBS have distinct effects on BEN in the striatum. Specifically, subthreshold stimulation led to a reduction in striatal BEN, while suprathreshold stimulation increased striatal BEN. This observation aligns with previous research using FC analysis on the same dataset, which demonstrated that subthreshold iTBS enhanced FC between the L-DLPFC and striatum. These differing effects of iTBS may induce varying projection patterns from the L-DLPFC to distant brain regions and increase top-down regulation of L-DLPFC to projection brain regions, particularly the striatum, which could be a critical brain region sensitive to iTBS intensities. We also found a close relationship between changes in BEN patterns induced by iTBS at varying intensities and the widespread distribution patterns of various neurotransmitter receptor and transporter binding maps. This suggests that the intensity of iTBS modulates the levels of multiple neurotransmitters, potentially elucidating its therapeutic role in various mental disorders. Additionally, our findings shed light on the neural mechanisms underlying different stimulation intensities and the diverse effects of iTBS.

### 4.1 No BEN changes in the stimulation target site

The effects of iTBS on BEN in the stimulation target site (L-DLPFC) were not observed with either subthreshold or suprathreshold stimulation. These results align with our previous findings from conventional HF-rTMS of L-DLPFC, where no alteration in BEN was observed in the L-DLPFC (Donghui Song, Zhang et al. 2017, Song, Chang et al. 2019). In our earlier study, the absence of change in BEN in the L-DLPFC may have been due to the timing gap between the rs-fMRI scan and HF-rTMS, potentially resulting in the effects of the L-DLPFC being mapped to distant brain regions rather than being observed directly at the target site (L-DLPFC). This phenomenon could also be relevant to the current study, as participants underwent a 7-minute break before the rs-fMRI scan. Furthermore, as in our previous studies, all participants in this study were healthy adults. The L-DLPFC plays a crucial role within the executive control network, overseeing multiple higher-order cognitive functions (Wagner, Maril et al. 2001, Friedman and Robbins 2022). In healthy individuals, higher-order brain functional networks typically exhibit relative stability. Given the robustness and intact normal functionality of L-DLPFC in healthy young adults, modulating brain activity within this region may pose challenges through single-session iTBS or HF-rTMS, especially when scanning was conducted several minutes after stimulation. It is noteworthy that there is still no consensus on whether TMS increases blood oxygen level-dependent (BOLD) activity at the site of stimulation (Bergmann, Varatheeswaran et al. 2021, Rafiei and Rahnev 2022). Recently a review from Farshad Rafiei et al (2022) suggested the absence of direct TMS-induced BOLD alterations at the stimulation sites and supported our findings that there is no change of BEN in the L-DLPFC. Current evidence suggests TMS induces periods of both heightened and diminished neuronal firing that counterbalance each other and consequently result in no net change in the overall BOLD signal after they reviewed all previous concurrent TMS-fMRI studies that reported analyses of BOLD activity (Rafiei and Rahnev 2022).

### 4.2 Reduced striatal BEN by subthreshold iTBS and increased striatal BEN by suprathreshold iTBS

The lack of change observed in L-DLPFC BEN after iTBS over L-DLPFC, along with the reduced striatal BEN by subthreshold iTBS, provides insights into the previously observed increase in frontal-striatum FC. This increased FC indicates enhanced neural synchronization between the striatum and the left DLPFC. However, it’s important to note that FC, defined by the correlation of time series, lacks a directional relationship. The findings from BEN suggest a potential directionality for this FC alteration: the enhanced synchronization may primarily result from changes in striatal neural activity rather than a shared change between the two regions. In other words, neural activity in the striatum appears to synchronize more closely with neural activity in the L-DLPFC, possibly due to top-down regulation from the L-DLPFC to the striatum.

As previously noted, L-DLPFC, a higher-order brain region responsible for executive functions, likely reflects its role in top-down regulatory functions, so the reduced striatal BEN may stem from this top-down regulation of L-DLPFC-striatum, as the frontostriatal circuit in modulating reward processing and cognitive control has been demonstrated in numerous studies and DLPFC projects to the caudate nucleus (Tekin and Cummings 2002, Chudasama and Robbins 2006, Robbins 2007, Staudinger, Erk et al. 2011, Morris, Kundu et al. 2016, Becker, Kirsch et al. 2017). In a clinical context, studies suggest that iTBS exerts top-down regulatory effects on the DLPFC-striatal circuit. A study has shown that smoking-induced craving relief relates to increased DLPFC-striatal coupling in nicotine-dependent women (Franklin, Jagannathan et al. 2021) and the relationship between the fiber connectivity integrity of the L-DLPFC-caudate and smoking cue-induced caudate activation can be mediated by the functional coupling strength of this circuit in smokers (Yuan, Yu et al. 2017). More importantly, subthreshold iTBS over L-DLPFC shows the beneficial effects in attenuating craving for cocaine, reducing intake, and prolonging abstinence in treatment-seekers (Sanna, Fattore et al. 2019), and FC of the L-DLPFC to striatum predicts treatment response of depression to TMS (Avissar, Powell et al. 2017). These studies underscore the modulatory effects of subthreshold iTBS over the DLPFC on the DLPFC-striatal circuitry. Indeed, this top-down regulation may specifically trigger striatal dopamine release. Recently, Shaikh et al. (2024) demonstrated through PET that subthreshold iTBS over L-DLPFC induces striatal dopamine release (Shaikh, Pellicano et al. 2024, Shaikh, Pellicano et al. 2024). Striatal dopamine is critical for many vital processes, including motivation, motor learning, and reinforcement learning, striatal dopamine supports reward expectation and learning (Calabro, Montez et al. 2023). Decreased BEN in the striatum suggests a greater tendency towards temporal consistency in striatal brain activity, then the striatal activity becomes more predictable and leads to increased long-range temporal coherence (LRTC) in the striatum (Wang 2021). This shows that subthreshold iTBS stimulation leads to a striatum signal containing more long-range memory during rest and then it may be more efficient at online information processing during tasks (He 2011, Liebana Garcia, Laffere et al. 2023).

We found an interesting result: subthreshold iTBS led to decreased striatal BEN, while suprathreshold iTBS resulted in increased striatal BEN. Corresponding to this intriguing finding, some studies indicate that both subthreshold and suprathreshold iTBS over L-DLPFC can ameliorate depression (Duprat, Desmyter et al. 2016, Blumberger, Vila-Rodriguez et al. 2018, Fitzgerald, Chen et al. 2020, Lee, Corlier et al. 2021, Plewnia, Brendel et al. 2021, Bulteau, Laurin et al. 2022). However, few studies have investigated the disparities between subthreshold and suprathreshold iTBS over L-DLPFC. In a study by Alkhasli et al (2019), homeostatic mechanisms were used to explain that the lack of an increase of fronto-striatal FC of suprathreshold iTBS, decreased activation or connectivity above a certain threshold and stronger stimulation could induce stronger surround inhibition at the stimulation side in the L-DLPFC and connected areas (Alkhasli, Sakreida et al. 2019). Another study by Sung Wook Chung et al (2018) found an inverse ULJshape relationship between intensity (50, 75, or 100% of rMT) and iTBS plastic effects and 75% iTBS yielded the largest neurophysiological changes, indicating that the assumption that higher intensity results in greater neuromodulatory effects may be false, at least in healthy individuals (Chung, Rogasch et al. 2018). In a clinical context, a current study indicates that subthreshold iTBS is associated with greater clinical efficacy than suprathreshold iTBS for MDD (Lee, Corlier et al. 2021). Based on the varying therapeutic effects of different stimulation intensities on depression, and the significant or even opposite relationships observed in the frontostriatal circuit for FC and BEN, one possible explanation for the occurrence of similar effects despite different stimulation intensities is that varying intensities triggering the release of different neurotransmitters. As previously mentioned, subthreshold iTBS results in DA release in the striatum and mental disorders often involve multiple neurotransmitters, such as depression being associated with DA, 5HT, GABA, and Glu (Nutt 2008, Duman, Sanacora et al. 2019)

### 4.3 The relationship between iTBS-induced BEN patterns and neurotransmitters

We performed a spatial correlation analysis to examine the relationship between iTBS-induced BEN patterns and neurotransmitter receptor and transporter binding maps. Our results demonstrate strong correlations between the BEN change patterns triggered by iTBS and various neurotransmitter receptor and transporter binding maps. Additionally, we noted notable variations in the correlation patterns between iTBS-induced BEN patterns across different stimulation intensities and receptor and transporter binding maps. Specifically, subthreshold iTBS-induced BEN patterns are predominantly associated with 5HT, CB1, GABA, M1, Glu, and VAChT. On the other hand, suprathreshold iTBS-induced BEN patterns are primarily linked to D2, DAT, and VAChT. Furthermore, we observed significant differences in the strength of correlations between BEN patterns induced by different stimulation intensities and these neurotransmitter receptor and transporter binding maps.

Although iTBS-induced BEN patterns and neurotransmitters were not directly derived from actual changes in the neurotransmitters of the participants in the study, this relationship is still reliable. As mentioned earlier, a large body of studies has demonstrated that rTMS can induce changes in neurotransmitter levels. Additionally, our study further confirmed the sensitivity of BEN to rTMS. Particularly noteworthy in this study is the observed interaction between stimulation intensity and time point within the striatum and this interaction is also evident in FC of the frontal-striatal network (Alkhasli, Sakreida et al. 2019). This alteration in spatial correlation indirectly suggests that iTBS induces changes in neurotransmitter levels, although the specific regions and magnitudes of these changes remain to be fully understood. Furthermore, the significance of the correlation coefficient in our spatial correlation analysis and the differences in correlation between BEN patterns and neurotransmitters induced by different stimulation intensities are evident, underscoring the robustness of this correlation. Some studies directly measuring neurotransmitter changes induced by iTBS over L-DLPFC also partially support this potential relationship. Iwabuchi et al. (2017) reported that subthreshold iTBS over the L-DLPFC significantly dampened fronto-insular connectivity and reduced GABA/Glu and GABA/Glu had a significant mediating effect on iTBS-induced changes in DLPFC-to-right anterior insula connectivity (Iwabuchi, Raschke et al. 2017). Recent studies have also shown that subthreshold iTBS over the L-DLPFC leads to DA release in the striatum using PET (Shaikh, Pellicano et al. 2024, Shaikh, Pellicano et al. 2024). In a clinical context, suprathreshold iTBS over L-DLPFC increased cortical GABA and Glu in treatment-resistant depression (Spurny-Dworak, Godbersen et al. 2022).

These differing neurotransmitters all relate to learning, memory, reward, and emotion regulation (Riedel, Platt et al. 2003, Gonzalez-Burgos and Feria-Velasco 2008, Koob 2009, Akirav 2011, Hayes and Greenshaw 2011, Meneses and Liy-Salmeron 2012), suggesting that the variances induced by these neurotransmitters may be critical in influencing the effects of iTBS at different intensities. Depression is associated with changes in these neurotransmitters (Nutt 2008, Palazidou 2012, Duman, Sanacora et al. 2019), and the interaction between these neurotransmitters may be the reason why different stimulation intensities can have therapeutic effects on depression (Blumberger, Vila-Rodriguez et al. 2018).

## 5 Conclusion

Our research provides new insights into the neural mechanisms underlying iTBS, enhancing our understanding of this novel NIBS protocol. In this study, we showed that the BEN is not only responsive to conventional HF-rTMS but also sensitive to iTBS, effectively capturing variations in neural activity induced by different stimulation intensities. Additionally, through indirect inference, we conducted a preliminary investigation into the relationship between iTBS-induced BEN patterns and neurotransmitter receptor and transporter binding maps, suggesting that different stimulation intensities may result in different neurotransmitter changes. Our findings highlight the importance of iTBS stimulation intensity and lay the groundwork for future clinical investigations aimed at identifying stimulation intensities with greater therapeutic benefits.

## Acknowledgments

We thank Alkhasli, et al for releasing their dataset.

## Data and code availability

All raw data are available in the online repository of OpenNeuro ds001832 (https://openneuro.org/datasets/ds001832/versions/1.0.1).

All neurotransmitter maps can be found at https://github.com/netneurolab/hansen_receptors/tree/main/data/PET_nifti_images.

All unthresholded statistical maps and the masks used in the study are available fro m https://neurovault.org/collections/QYIWAEYS/.

For the preprocessing and analysis of images, fmriprep is available from https://fmriprep.org/en/0.6.4/index.html. Nilearn is available from https://nilearn.github.io/dev/index.html. SPM is available from https://www.fil.ion.ucl.ac.uk/spm/software/spm12/. AFNI is available from https://afni.nimh.nih.gov/.

Further updates related to this study will be available at https://github.com/donghui1119/Brain_Entropy_Project/tree/main/NIBS/ITBS (upon publication of the manuscript).

## CRediT authorship contribution statement

Pan-Shi Liu: data analysis, visualization, manuscript drafting, and editing. Dong-Hui Song: conceptualization, data analysis, visualization, manuscript drafting and edit ing, supervision, project administration. Xin-Ping Deng: manuscript drafting and editing, Yuan-Qi Shang: manuscript editing. Qiu Ge: manuscript editing. Ze Wang: conceptualization, manuscript editing and final manuscript proofing, supervision, project administration. Hui Zhang: manuscript editing, supervision, project administration, acq uisition of funding.

## Notes

https://openneuro.org/datasets/ds001832/versions/1.0.1

### Competing Interest Statement

The authors have declared no competing interest.

